# Recommendations for Automating Hydrogen/Deuterium Exchange Mass Spectrometry Measurements using Data-Independent Acquisition Methods

**DOI:** 10.1101/2025.01.18.633743

**Authors:** Frantisek Filandr, Morgan Hepburn, Vladimir Sarpe, D. Alex Crowder, Maryam Hassannia, Stephen Coales, Yuqi Shi, Rosa Viner, Martin A. Rossotti, Joey G. Sheff, Jamshid Tanha, David C. Schriemer

**Author notes:** Authors contributed equally (Frantisek Filandr, Morgan Hepburn, Vladimir Sarpe).

## Abstract

Hydrogen/deuterium exchange mass spectrometry (HX-MS) is a method used to study solution-phase protein structure and dynamics. Despite its many applications, HX-MS is limited in throughput because manual data analysis is still the norm. We previously developed HX-MS^2^ technology to add a second dimension of deuteration data and promote automated data processing. Data-independent acquisition (DIA) techniques enable this approach, but we require optimized methods for best performance. Using an Orbitrap Eclipse for illustration, we show that ion optics and collision energy settings typical of a proteomics DIA experiment generate maximal peptide retrieval from the DIA library. As few as three MS^2^ sequence ions are sufficient to generate a deuteration measurement with a precision that exceeds what is possible in traditional HX-MS. DIA window sizes are based on the chromatographic resolution of the method. An inter-scan window offset method is the recommended default configuration for most HX-DIA applications, but an intra-scan overlap method can be tuned for highest performance and is recommended when maximum peptide retrieval is desired. A fully automated HX-MS^2^ solution consists of Trajan HDX automation technology, an Orbitrap Eclipse mass spectrometer and AutoHX software. We demonstrate its robustness on an extensive time-course analysis of phosphorylase B and an epitope analysis of single domain antibodies (V_H_Hs, nanobodies) specific to the receptor binding domain of SARS-CoV2 spike protein.

## Introduction

Hydrogen/deuterium exchange mass spectrometry (HX-MS) is a structural mass spectrometry method capable of monitoring the structural dynamics of proteins in solution, under conditions close to their native state^1–3^. It measures the exchange of protein backbone amide hydrogens for deuterium when the protein is exposed to a D_2_O-containing buffer. Any changes to solvent accessibility and/or hydrogen bonding can result in a measurable alteration of deuterium uptake. The method supports a wide range of applications, such as the analysis of protein-protein and protein-drug interactions^4–8^, the monitoring the structural effects of mutations^9,10^, and the measurement of protein stability^11,12^. A standard peptide-based HX-MS method can usually localize binding sites and conformational adaptations to within a few amino acid residues.

The method still finds limited use unfortunately, even though new tools have improved accessibility^1^. The main issue preventing HX-MS from becoming a scalable assay technology is low throughput. Sensitivity has improved considerably with the use of proteomics-grade front-end systems^13^, but sample throughput is restricted by time-consuming manual data curation. An expert in the field is required to visually inspect the quality of peptide isotopic distributions and error-check software determinations. A typical two-state differential HX analysis of a single protein system can require two hours of mass spectrometer acquisition time and six to eight hours of manual data curation with current software. The time burden scales with the number of conditions analyzed, the number of timepoints in the analysis, and the size of the protein system. It is not unusual for more complex experiments to require months of manual data curation^14,15^. Large-scale experiments like drug candidate screening involving anything other than simple pure proteins are out of reach for most labs.

We previously demonstrated the advantages of adding a second dimension of deuteration data in the form of collision-induced dissociation (CID)-generated peptide fragments^16^. CID using beam-style ion activation events scrambles deuteration throughout the peptide, rendering each sequence ion a surrogate peptide measurement. These fragments overcome the information deficit that forces the manual inspection of MS-only data. We demonstrated that fragment deuteration can be used to authenticate peptide selections, generate deuteration measurements, and avoid the MS measurement altogether in cases of overlapping isotopic distributions in MS^1^ data. There is no increase in resolution with CID-based HX-MS^2^ as there is in Electron-Transfer Dissociation (ETD)-based HX-MS^2^, but it can deliver peptide measurements with accuracy and precision. At the time, data-dependent acquisition (DDA) methods were not effective for reliably selecting deuterated peptides for fragmentation analysis, and Data Independent Acquisition (DIA) technology was in its infancy. Initial applications of the approach, while successful, were not very easy to implement^17,18^.

Newer mass spectrometers provide windowed DIA methods that can collect rich peptide-fragment deuteration data for all peptides entering the instrument^19^. These improvements spurred us to develop AutoHX software and exploit the extra dimension of data^20^. To accurately measure the deuterium level for each peptide, the software generates a linear deuteration model using the fragment and precursor deuteration data. Deuteration values calculated from the models are demonstrably better than those obtained by MS-only data analysis, and results are returned much faster^20^. The software is fully automated and scalable to virtually any size of experiment, requiring processing times of only minutes per file.

Here we describe how to create a windowed DIA-based HX-MS^2^ experiment suitable for the analysis of simple or complex protein samples. We also describe additional improvements to AutoHX software that simplify the production of high-quality and reproducible analyses. The study is based upon an Orbitrap Eclipse mass spectrometer, but we highlight principles that can be extended to any instrument that can make windowed DIA measurements.

## Methods

### HX-MS^2^ of Phosphorylase B

Sample handling was completed with Trajan HDX automation. Phosphorylase B (Sigma Aldrich) was resuspended in HEPES buffer 1 (25mM HEPES, 150mM NaCl, pH 7.2), to final concentrations of 2.3 and 5.7 µM, for injections of 50-100 pmol. For mapping experiments, phosphorylase B solutions were diluted 1:1 (v/v) with HEPES buffer to emulate labelling, or 1:1 with D_2_O buffer (25mM HEPES, 150mM NaCl pD 7.2 in 100% D_2_O), for HX experiments. Samples were then mixed 1:1 (v/v) with quench buffer (500 mM Glycine HCl, pH 2.3, 6M Urea) and injected for liquid chromatography (LC)-MS analysis on a Vanquish Neo-HDX LC coupled to an Orbitrap Eclipse (Thermo Fisher Scientific). Samples were digested online using a Nepenthesin-2 column (Affipro, CZ), loading at 110-170 µL/min of mobile phase A (0.4% formic acid). Loading/digestion flow rates were optimized as needed. Peptides were trapped and washed on an Acclaim trap cartridge (1mm x 5mm, Thermo Fisher Scientific) before separation on a Hypersil GOLD C18 column (1mm x 50mm, Thermo Fisher Scientific) at 40 µL/min, using a 10-17 min gradient of 4-45% mobile phase B (80% acetonitrile, 0.4% formic acid).

For peptide library generation, data were collected in DDA mode, with MS^1^ acquired over m/z 375-1000, 60K resolution, a radio frequency (RF) setting of 40, and AGC at 100%. Peptides were fragmented with higher-energy collisional dissociation (HCD) at 30% normalized collision energy (nCE). MS^2^ was set with an automatic gain control (AGC) of 300% and a maximum injection time of 54 ms, at 30K resolution. For labelled samples, data were collected in DIA mode, with MS^1^ acquired over m/z 375-1000 at a resolution of 60K. DIA MS^2^ scans were acquired at a resolution of 30K, AGC of 300%, a maximum injection time of 54 ms, and the respective parameters were varied per experiment. In the RF, ion transfer tube (ITT) temperature, and nCE optimizations, we used an intra-scan method where the DIA windows were 20 Th wide with a 2 Th overlap. For the DIA window width experiments, a 2 Th overlap was used for each window size, whereas a 10 Th window was used for the window overlap experiment. A 20 Th window was used for the inter-scan method. Unless otherwise indicated, the HCD nCE was set to 25%, the RF to 35 and the ITT at 320 °C.

All data were processed in Mass Spec Studio 2.0 (version 2.4.0.3661). Mapping runs were searched against phosphorylase B and Nepenthesin-2, using the HX-PIPE app in the Studio^21^, which is based upon MS-GF+ for peptide identification^22^. MS^1^ and MS^2^ error tolerances were set at 10 ppm, digestion as unspecific, charge states as +1-6, with a minimum peptide length of 5, and peak to background ratio set to 0. Peptides with a p-value less than 0.05 were accepted and exported, merging duplicate peptide identifications within +/-0.4 min into a single peptide identification. The AutoHX app was then used to quantify deuterium uptake and determine the number of valid peptides. Basic parameters related to LC and MS instrument performance were specified as follows: MS^1^/MS^2^ mass tolerances were set to +/-10 ppm with a peak to background ratio of 0 (for non-Orbitrap data this should be adjusted higher), extraction ion chromatogram (XIC) integration range was set to 0.2 min with an allowable retention time (RT) variance of ±0.4 min, and smoothing turned off. (This smoothing has no effect on data processing as is applied purely for visual inspection; leaving it off allows the user to better inspect the chromatographic sampling rate). In experiments with 3 replicates per state, the “% minimum valid replicates per set” was 100% (all three replicates must be used to be considered valid), but when using four replicates per state it was set to 75% (for three out of four replicates). All default processing parameters were left unchanged. In cases of low sequence coverage, the user can turn on “Accept MS1 with Minimal Qualifier Sequence Ions”, which allows the rescue of high-quality peptides that are identified in the map but did not fragment well in DIA.

Once the Auto-HX processing is complete, a results summary tab displays information on global data quality and provides opportunities to refine the analysis, primarily by varying the minimum number of quantifiable fragments and the Deuteration Error Filter (Max %D standard deviation). The summary tab helps inspect the balance between sequence coverage and noise when we vary these two parameters. The default minimum of 3 fragments and 5% error can be too permissive at times. A range of 3-6 fragments and a maximum 3% standard deviation was used for most experiments presented, except where noted. For differential analyses, the State Comparison tab was also useful when refining these two parameters.

### HX-MS^2^ of Nanobodies

V_H_Hs were expressed in *Escherichia coli*, and RBD/SD1 was expressed in HEK293-6E cells and purified as described previously^23^. Sample handling was completed with Trajan HDX automation. The RBD/SD1 from the ancestral SARS-CoV2 spike protein (aa319-591) and the corresponding nanobodies were diluted in HEPES buffer 2 (25 mM HEPES 150 mM NaCl, pH 7.4) to final concentrations of 6 µM, for a 50 pmol injection. The RBD and nanobody solutions were combined 1:1 (v/v) and incubated for 1 min to allow for equilibration of the antigen/antibody complex. The unbound RBD state was diluted 1:1 with HEPES buffer 2 in the absence of nanobody. For mapping, complexes were diluted 1:1 (v/v) with HEPES buffer to emulate labelling, or 1:1 (v/v) with D_2_O buffer 2 (25 mM HEPES, 150mM NaCl pD 7.4 in 100% D_2_O) for HX experiments, and labelled for 1 min. The samples were then mixed 1:1 (v/v) with quench buffer (500 mM Glycine HCl pH 2.3 6M Urea) for digestion and LC-MS analysis on a Vanquish Neo-HDX coupled to an Orbitrap Eclipse, as described above, except that separation occurred over an 8 min gradient of 4-45% mobile phase B. Mapping runs were collected as above. The HX samples were also collected as previously described but using an nCE of 26% and applying the intra-scan DIA method with a 10 Th window and a 2 Th overlap. Data were processed on Mass Spec Studio 2.0 mostly as above, searching the mapping files against the RBD and Nepenthesin-2 sequences. AutoHX was used for differential analysis, defining unbound RBD as the control to auto-populate the control state in results comparison tab, and to support the Woods plots. The minimum number of fragments was set to 5 and the maximum % D standard deviation was set to 3. Peptides with significant changes in deuteration were defined based on the Woods plots, using overlapping peptides to estimate higher resolution where justified. Significant changes were mapped to the SARS-CoV-2 spike structure (PDB 6VXX)^24^ using ChimeraX, with residues not in the partial RBD removed.

## Results and Discussion

### Defining the Peptide List

The DIA-based HX-MS^2^ experiment requires a sequence map of the protein(s) being studied, as is typical of the conventional HX-MS method. The map is produced by running DDA-based analyses of unlabeled protein digests and using an appropriate search tool on the data. We used HX-PIPE, the internal search engine in the Mass Spec Studio^21^, to identify peptides in the digest of phosphorylase B. Other database search engines can be effective as well, provided they are designed to search digests arising from non-specific enzymes. The full list of peptides defines the peptide library (see reference 21 for the file format), which is then fed directly into AutoHX along with the DIA datasets collected on the deuterated samples.

The goal of a DIA-based HX-MS^2^ experiment is to maximize the number of peptides from the library that can be used as reliable reporters of deuteration, which we define as *recall*. Ideally all peptides in the library would be recalled. Any fragments identified during the database search should support deuteration measurements, however the quality of fragment isotopic distributions will vary. A successfully recalled peptide must pass a strict set of criteria. First, basic signal quality standards for the precursor and the associated fragments must be met or exceeded (**Figure 1A**). Second, a linear deuteration model must be built from a minimum number of high-quality fragment ions (**Figure 1B**). Only fragments within a chosen model precision are accepted. The rest are designated as outliers and not used for deuteration calculations (grey dots in the figure). An optimization process is conducted that chooses a subset of the inlier fragments to define the model (red dots in the figure). The precursor can be part of this calculation. The model can exclude some acceptable fragments (black dots in the figure) if the optimization process finds a higher precision subset of fragments.

**Figure 1.**
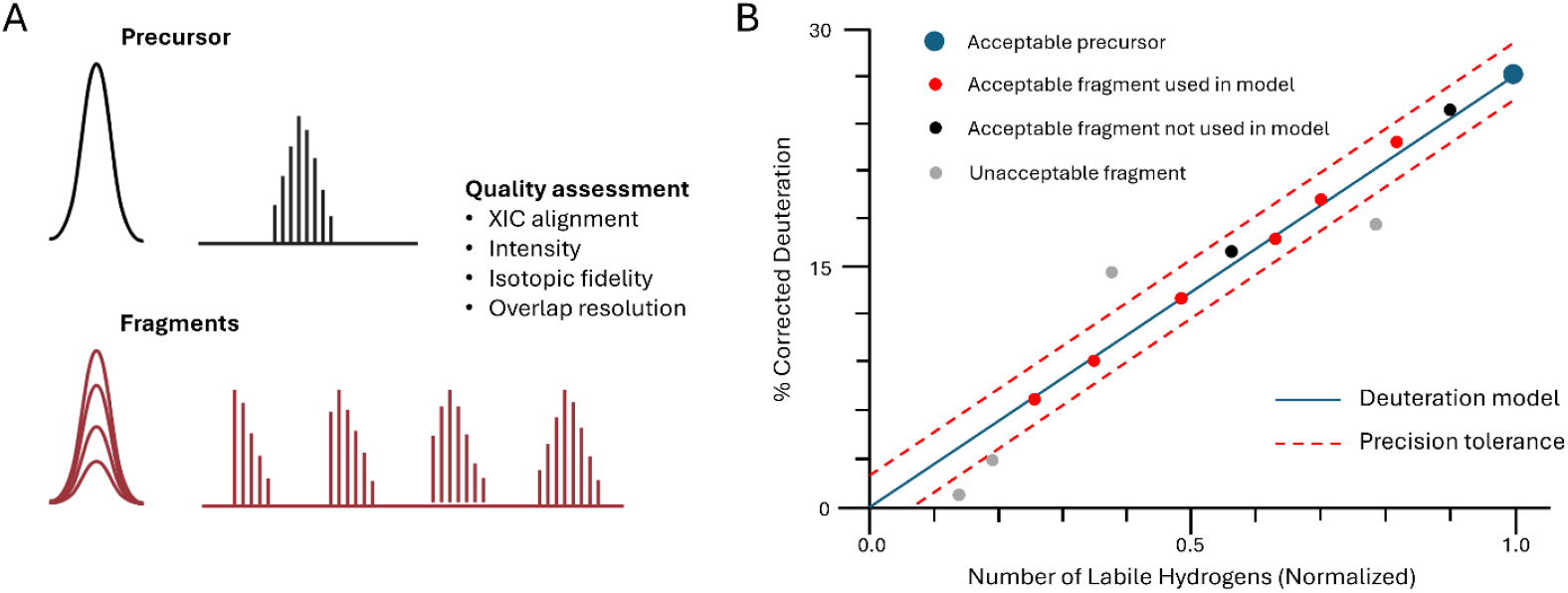
Key elements of an HX-DIA experiment. (A) Deuteration analysis is conducted in both the MS^1^ and MS^2^ domains, on isotopic distributions arising from aligned XICs that pass minimal quality standards. (B) Linear deuteration model generated from the full set of precursor and fragment isotopic distributions. The model is generated from only the highest quality subset of distributions and is used to calculate a consensus peptide deuteration value across all replicates.

We specify two classes of fragments: quantifiers and qualifiers. Quantifiers are fragments that can be used to generate the model (red and black dots). Qualifiers establish the identity of the peptide. They include the quantifiers and any model outliers that meet a minimum spectral quality (some of the grey dots). Thus, a recalled peptide is one that generates enough quality fragment information to confirm peptide identity in the deuteration experiment, and one that produces a deuteration model defined by a minimum set of quantifier fragments. The subset of fragments is chosen to maximize precision across replicate measurements. Together, this process delivers the best and most reliable per-peptide data for the deuteration calculation.

Fragments dominate the measurement. There is only one instance in which precursors are used exclusively. Certain peptides will not generate a rich fragment spectrum but will produce a good MS^1^ spectrum (*e*.*g*., a small singly charged peptide). In this case, the precursor can be used to define the deuteration model provided that the MS^1^ spectral quality is very high, and qualifier fragments are present to license the selection.

### Maximizing the Number of Peptide Reporters

With these criteria in hand, we explored the effect of major variables in an HX-DIA experiment on peptide recall, using phosphorylase B as our test case. We first evaluated if non-standard ion optics settings might be necessary as the method relies upon deuterium scrambling^16^ and the deuteration models are slightly biased towards larger sequence ions^17^. The RF lens (**Figure 2A**) and ion transfer tube settings (**Figure 2B**) had little effect upon the number of recalled peptides. The optimal collision energy is comparable to a typical proteomics experiment, and it varies by only 25% across a wide range (**Figure 2C**). Thus, there is considerable latitude in establishing ion desolvation and transmission conditions for a given experiment.

**Figure 2.**
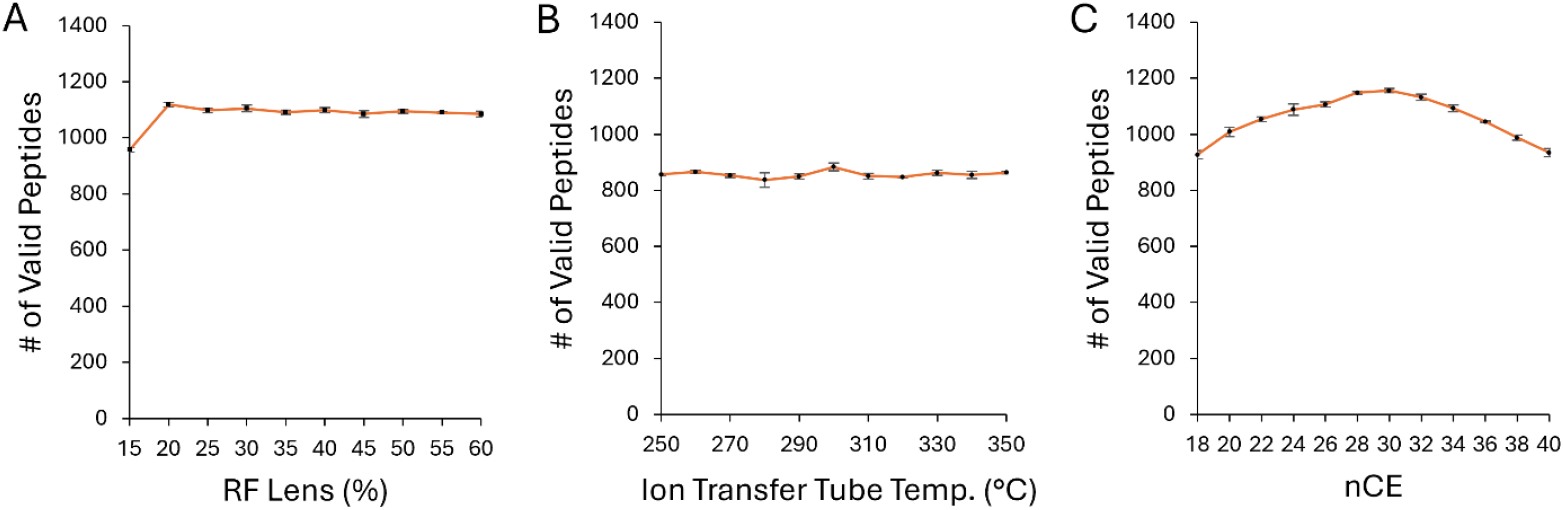
Effect of basic ion optics settings on the number of valid peptides recalled from the library. (A) RF setting. (B) Temperature of the ion transfer tube used in desolvation. (C) Normalized Collision Energy (nCE). The samples in B used a lower injection amount compared to A and C. Error bars represent +/-one standard deviation (n=3 replicates).

### Optimizing Sequence Coverage vs. Measurement Precision

AutoHX allows the user to specify a minimum number of fragment ions for selecting a peptide. This variable is the user’s primary tool to mine the peptide library. To investigate the relationship between peptide recall and the minimum number of required fragments, we processed a replicate set of phosphorylase B data for a given labeling timepoint (3 min). Not surprisingly, the number of validated peptides dropped significantly as a function of fragment number (**Figure 3**) and the average peptide length increased (**Figure S1**).

**Figure 3.**
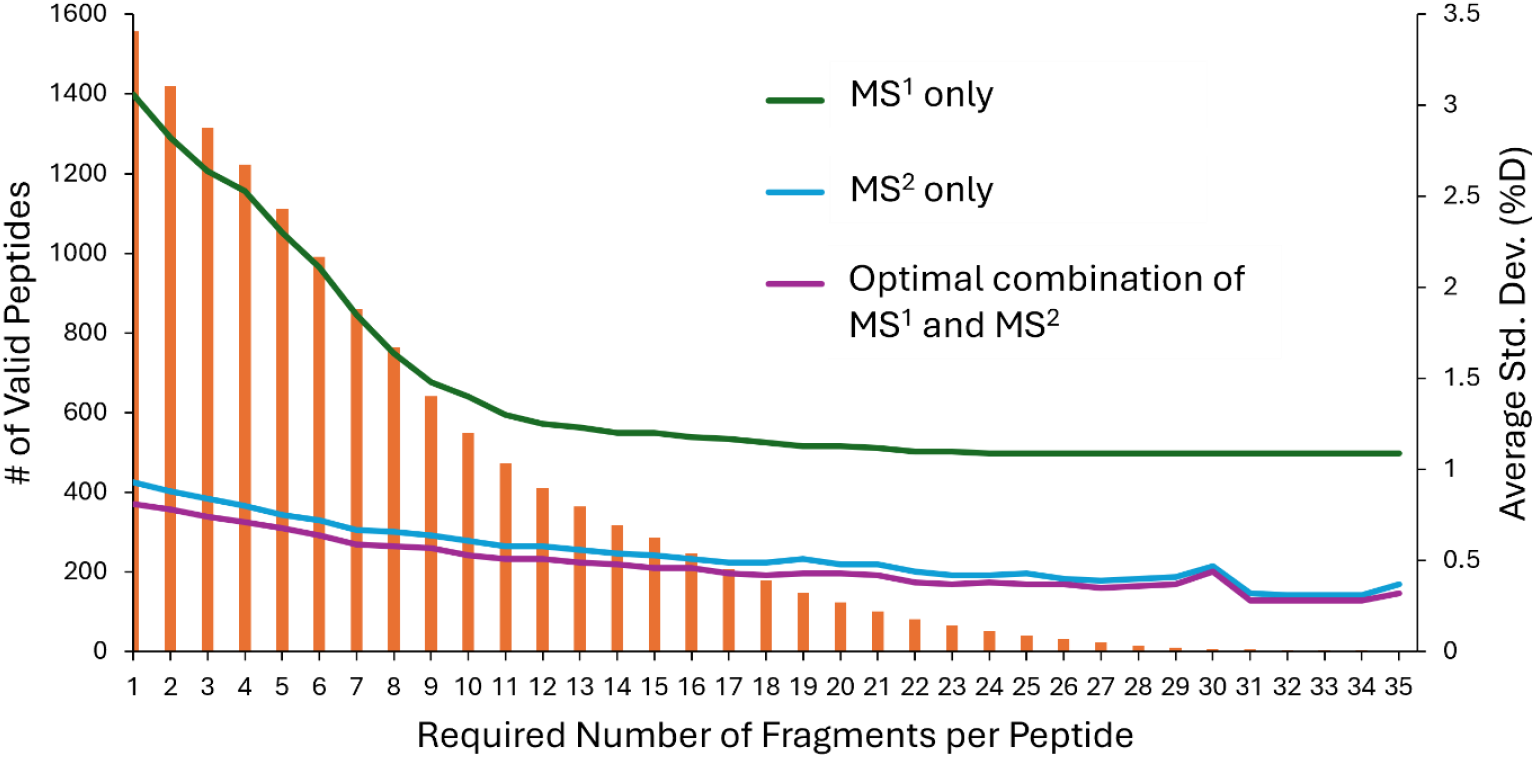
Impact of fragment selection on peptide recall and measurement precision. The number of valid peptides recalled from the library (orange bars) is shown as a function of the minimum number of quantifier fragments required to validate a peptide. The corresponding average standard deviation (Std. Dev.) of the deuteration measurements (n=3 for each peptide) displayed three ways: (i) using the data from the precursor alone (green line), (ii) the data from the fragments alone (blue line), and (iii) the best combination of the two (purple line).

The effect of fragment number on precision is more interesting. DIA methods generate both MS^1^ and MS^2^ spectra, which support alternative calculations of deuteration levels and precision. We can just use the MS^1^ deuteration data as in a conventional measurement or use fragment deuteration data, with or without the precursor data added in. The precision of MS^1^- only measurements is strongly dependent on the minimum number of fragments. Those based solely on MS^2^ measurements are weakly dependent on the minimum number of fragments. This is sensible. In general, peptides that produce fewer useful fragments are of lower abundance and generate a lower signal to noise ratio (S/N) in MS^1^. A lower S/N will impact the sampling of the isotopic distribution and reduce precision. However, the MS^2^ dimension improves S/N by reducing noise. Further, the software can choose the highest quality and most precise subset of fragments to support the deuteration measurement. This selection process preserves high precision across a wide dynamic range. Allowing the algorithm to sample MS^1^ values together with the MS^2^ values increases the precision further, although only by a small amount (**Figure 3**).

Taken together, the user has considerable flexibility in setting the minimum number of quantifying fragments required and can modify the value as needed to maximize sequence coverage and redundancy. It should be noted that error rates can increase based on misidentifications, thus very low fragment numbers should be avoided in complex protein mixtures. We recommend a minimum of three fragments and routinely use three to eight. Note that these optimizations were implemented using the intra-scan DIA method with a window size of 20 Th and an overlap of 2 Th, as described below.

### Optimizing the DIA Experiment – Window Size and Overlap Strategies

DIA methods specify a transmission window size (the “window”) and usually an overlap between adjacent windows to faithfully sample the entire, undistorted set of isotopic distributions^17^. Window widths are established empirically, balancing the chromatographic peak widths, the complexity of the sample, and the effective sweep rate of the instrument. Standard proteomics experiments often use a 1 Th overlap between adjacent windows so that most of the isotopic envelope will exist in one or the other window. It needs to be larger in an HX-DIA experiment because the isotopic envelope expands with deuteration. There are multiple strategies for creating a DIA experiment, depending on the instrument used^19^, and most will work with AutoHX. For the Orbitrap Eclipse, we can specify intra-scan DIA or inter-scan DIA. The intra-scan method creates an overlap between adjacent windows as described above (and in **Figure 4A**), whereas the inter-scan technically has no overlaps, but rather an offset between two adjacent scans of the m/z range (**Figure 4B**). We first explored the intra-scan method. We fixed the overlap at 2 Th and then measured recall as a function of window width, from 5 – 60 Th (**Figure 4C**). The number of valid peptides recalled was maximal at 10 Th and then decreased slowly. However, edging was quite high when using a 5 – 10 Th window. Edging is declared (and reported by the software) when the transmission window may distort the isotopic distribution of a deuterated peptide, based on the proximity of the leading or trailing isotopologues to a window edge. We specified a proximity of 0.5 Th on the left and 1.0 Th on the right as the edging criteria for the Eclipse. Larger windows will obviously generate fewer potential edging events in a scan because there are fewer windows per scan, but these larger windows create more complex MS^2^ spectra, which in turn has a negative effect on fragment detection and thus recall. These effects counteract each other, hence the optimum at 10 Th.

**Figure 4.**
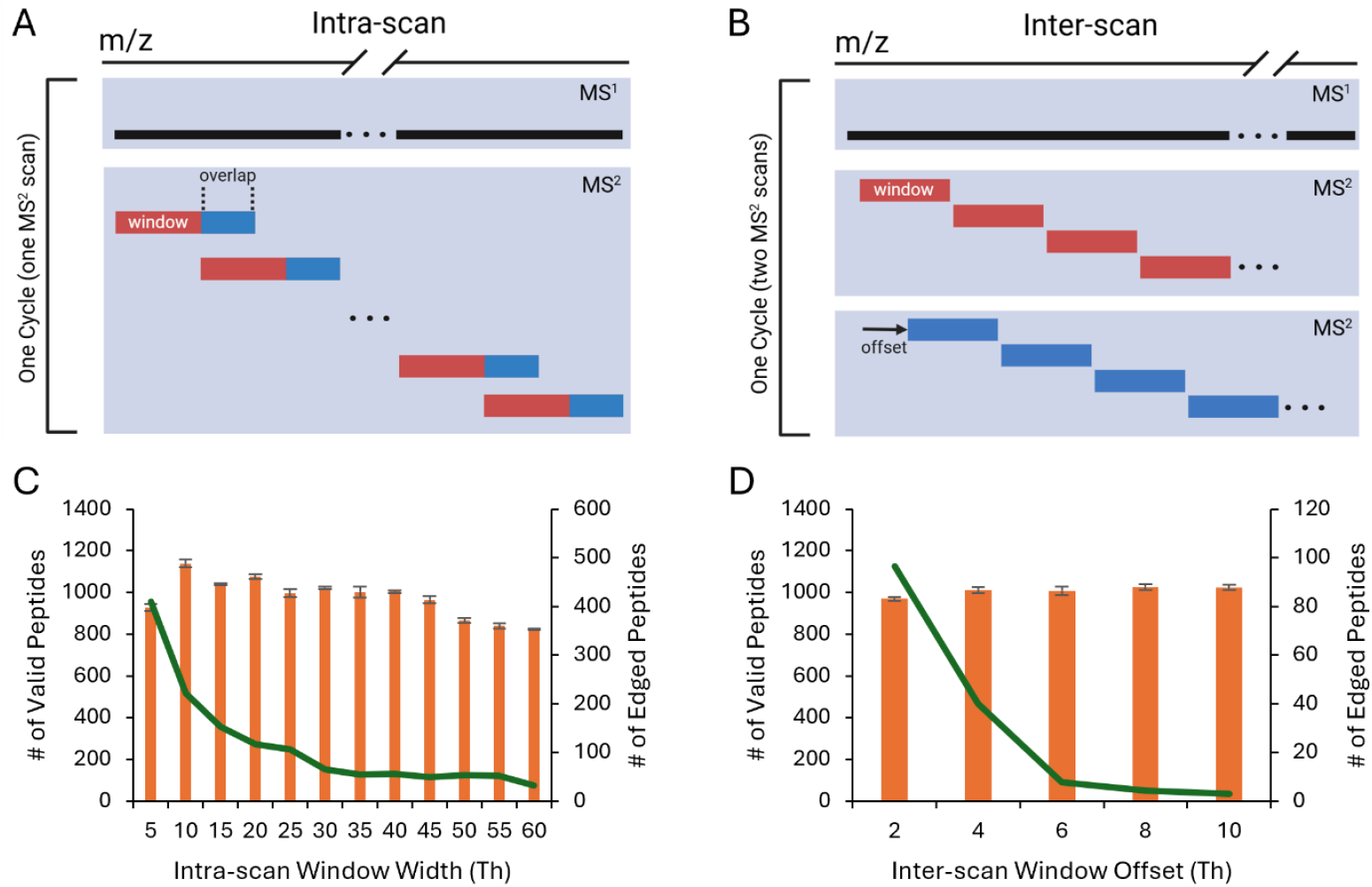
DIA strategies for collecting HX data. (A) Representation of the intra-scan method, where the MS^1^ acquisition is paired with one scan of the m/z range for MS^2^ acquisition, using overlapping windows. Both window size and overlap are user-specified. (B) Representation of the inter-scan method, where the MS^1^ acquisition is paired with two scans of the m/z range for MS^2^ acquisition, implementing an offset between scans. Both window size and offset are user-specified. (C) Effect of the intra-scan window width on the number of valid peptides recalled from the library (orange bars) and the number of peptides that are edged, or potentially compromised, by window placement (green line). The window width represents the value input into the instrument’s method editor; the true window width equals the input value plus the overlap (here 2 Th). Data from the 3 min timepoint of a phosphorylase B labeling experiment (50% D_2_O) (D) Effect of the inter-scan window offset using a 20 Th window; graph details as in C. Error bars represent ±1 Std. Dev. (n=3). Data from the 5 min timepoint of a phosphorylase B labeling experiment (50% D_2_O).

We then fixed the window size at 10 Th and varied the overlap from 0 to 5 Th. The number of edges remain the same over this range, but the wider overlaps mean that more peptides will be fully sampled in at least one of the adjacent windows. Edging is effectively abolished at 5 Th but an overlap as low as 2.5 Th already generates maximum recall, which levels out as the overlap is increased (**Figure S2**). That is, although edging is steadily reduced from 2.5 – 5 Th, no additional peptides are recalled. There are a few reasons for this leveling out. First, the effective window size increases with the overlap, which we showed can counteract an increase in recall (**Figure 4C**). Second, our edging definition is quite permissive so the software can already rescue some peptides that approach the edges, and third, if an edged peptide is of low quality to begin with, it cannot be recalled. We recommend reducing edging as much as possible using a combination of larger overlaps (to minimize the effects of edging) and larger windows (to decrease the number of edges).

Taken together, for the intra-scan method on a tribrid like the Eclipse, we recommend using a window in the range of 10 – 20 Th. An overlap of 2 – 5 Th should be applied when using 50% or less D_2_O in the labeling experiment, or 5 – 10 Th overlap when using 75-90% D_2_O. The choice of the window size should always be made with reference to the chromatographic peak width of the method to ensure adequate sampling (**Figure 5A**). We seek to sample a chromatographic peak approximately five times. A 20 Th window will perform well when the minimum chromatographic peak widths are 6 sec (full width half maximum, FWHM) and the scan is from m/z 375-1000. Note that cycle times are not dependent upon the size of the overlap, so it can be varied as needed.

**Figure 5.**
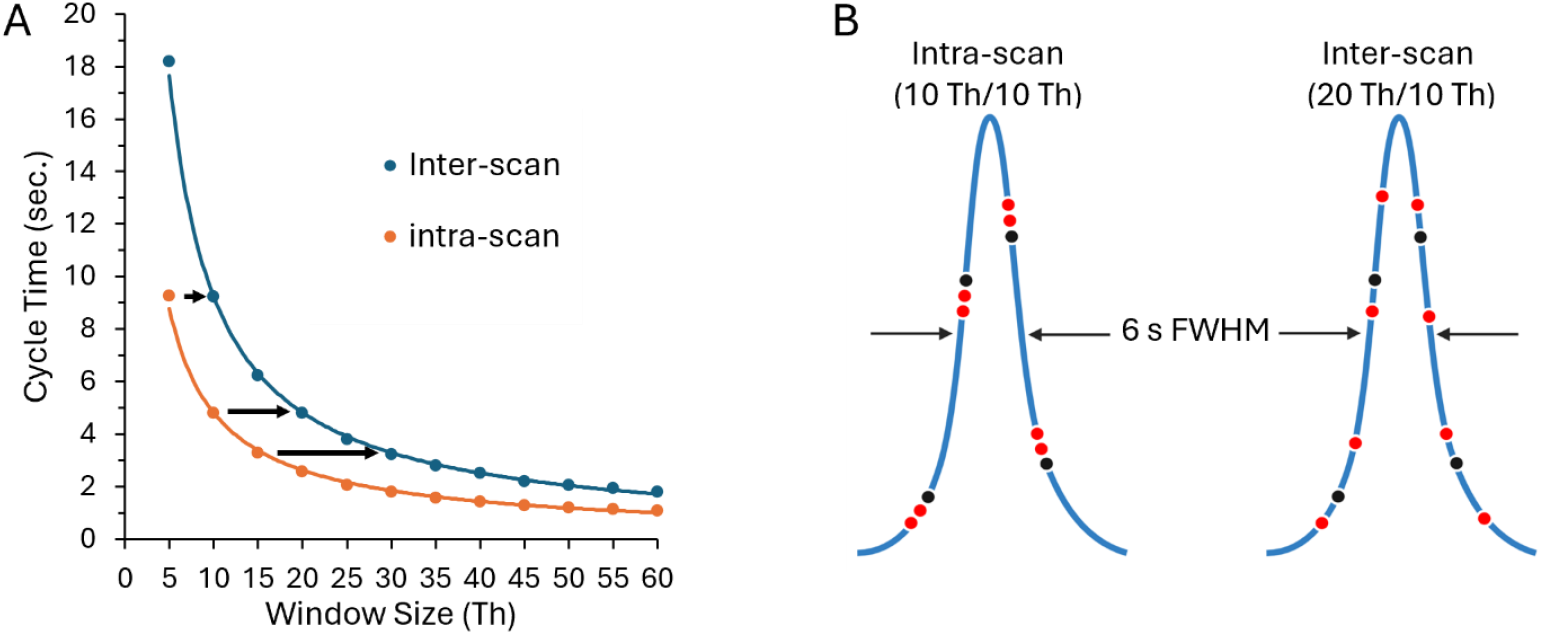
Chromatographic sampling dependency on window size and placement for the Orbitrap Eclipse. (A) The dependency of MS cycle time on window size and DIA method, measured on an Orbitrap Eclipse, averaged from data presented in Figure 4. When the overlap for the intra-scan type approaches 100% of its window width then the effective window width (highlighted by black arrows) approaches the inter-scan method. (B) The chromatographic sampling rate doubles for peptides that are never edged (red dots); edged peptides retain the sampling rate based on figure A (black dots). The double sampling for the intra-scan method is closely spaced but more dispersed for the inter-scan method. The figure is modeled based upon the sampling of a chromatographic peak-width of 6 sec FWHM and 4.8 s per point, selected from figure A, with the specified window/overlap size.

An interesting advantage occurs when the chosen overlap approaches 100% of the window size. The entire m/z range is covered twice per scan, effectively doubling the sampling rate for a given cycle time, at least for all the peptides that do not experience edging. At this point the second DIA strategy can be more effective (**Figure 4B**). The inter-scan method requires two MS^2^ scans per cycle to solve for edging, which can easily be eliminated with an offset of 6 Th or higher (**Figure 4D**). The basic sampling rates of the two methods are identical when the window in the inter-scan method is double that of the intra-scan method (**Figure 5A**). However, note that the effective window sizes are the same when the intra-scan’s overlap is 100% of its window size. For example, for the inter-scan method, a 20 Th window swept twice with a 10 Th offset generates the same effective window size as an intra-scan method with 10 Th window and a 10 Th overlap, and the cycle time is 4.8 sec for both on the Eclipse. The inter-scan method always doubles the sampling rate for peptides that are not edged, and it does so more effectively than the intra-scan method. It samples chromatographic peaks more evenly because the second sampling occurs in the next scan, rather than the next window (**Figure 5B)**. Regular sampling is preferred for better chromatographic peak definition and improved performance overall. Of course, not all peptides will benefit from this 2x sampling rate. In the phosphorylase B inter-scan DIA experiments, between 50-60% of the peptides were double sampled with a 20 Th window (not shown). The remainder will be sampled at the basic rate (**Figure 5B**).

Taken together, we recommend choosing the inter-scan method as the default HX-DIA strategy and then adapting as required. For chromatographic peak widths of 6-10 sec FWHM, we suggest selecting a 20 Th window and a 10 Th offset for the Eclipse or similar instrument. This recommendation is independent of the %D_2_O used in labeling. A 2:1 ratio of window to offset should always be preserved to take full advantage of the 2x sampling rate for peptides that are never edged, and the actual values scaled higher or lower depending on instrument scan speed and chromatographic resolution. The inter-scan method is robust and will be ideal for most applications. We use the intra-scan method, and the recommendations we made above, when the complexity of the sample is high (*i*.*e*., multiple proteins, contaminated proteins, or very large protein complexes) or when the chromatographic resolution is 6 sec FWHM or better. Here we typically use an overlap less than half of the window width and reduce labeling to 50% D_2_O or lower when needed^25^. The intra-scan method offers some tunability, which can be useful when attempting to improve coverage in select regions of a protein sequence. Finally, note that the AGC setting must also be considered when selecting a window width. Compared to small windows, large windows can reduce fragment intensity by filling the trap much more rapidly (**Figure S3**), which could negatively affect fragment selection. We recommend a setting of 300% AGC for a total window width of 20 Th (window plus overlap for the intra-scan method); higher AGC limits could be explored for larger windows and/or complex protein digests.

### Automated Application #1 – Extensive Kinetics

To illustrate the robustness of the complete package, we performed an extensive kinetics analysis of phosphorylase B. We used Trajan HDX automation to generate 22 timepoints in quadruplicate resulting in 88 samples in total. Data acquisition required 36 h. We specified a minimum of 6 quantifiable fragments per peptide in AutoHX, which selected 825 common peptides across all timepoints in at least 3 out of 4 replicates, resulting in 98.8% sequence coverage and a 25.1-fold redundancy, with an average peptide length of 15.7 amino acids. Automated data processing on a desktop computer required 4.5 h, one-eighth of the acquisition time, which included the generation of visual reports such as a summarizing heatmap and sequence map (**Figure 6, Figure S4**). No outliers are apparent and both fast and slow exchanging regions are clearly visible along the protein sequence. The dataset contained 187,880 individual raw peptide deuteration measurements. Conventional manual validation is completely impractical for this exercise. If we presume five seconds to manually validate one peptide measurement, over 32 eight-hour workdays of non-stop effort would be required to curate such a dataset.

**Figure 6.**
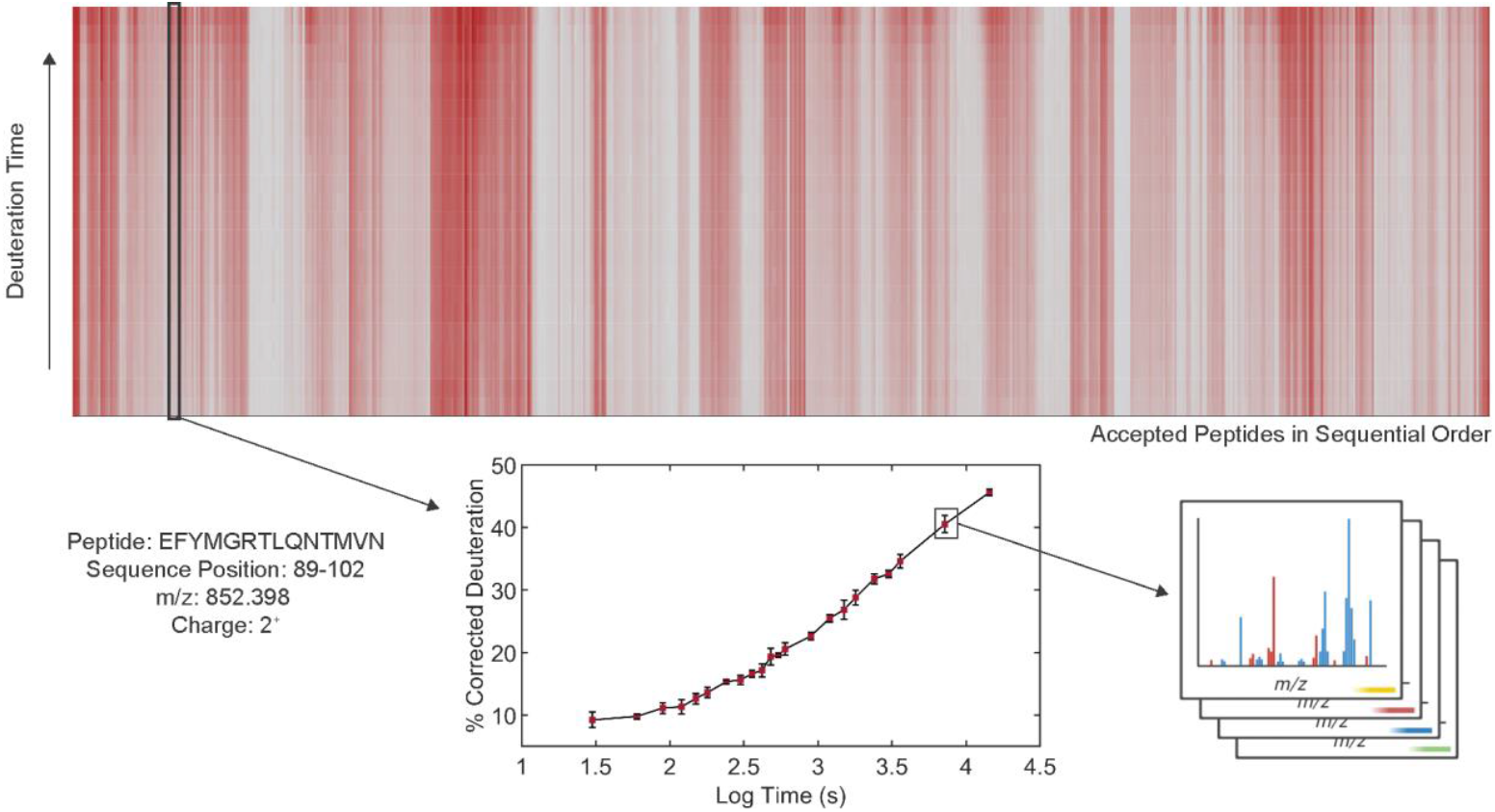
Kinetics analysis of phosphorylase B. A 22-point time course was collected, with each time point collected in quadruplicate. The data were collected with the intra-scan method using a window of 10 Th and an overlap of 2 Th and applying a minimum requirement of 6 fragments. All defaults were used in the AutoHX software, except for “peak to background” (=0) and the “minimum number of replicates” (=75%). Heatmap displays labeling by all 22 deuteration times (vertical) and all 825 peptides ordered according to position in sequence. Inset shows a kinetics trace for a randomly selected deuterated peptide. Error bars represent ±1 Std. Dev. (n=3,4).

### Automated Application #2 – Differential Analysis

Protein or ligand binding experiments are the most common applications of HX-MS, designed to map binding sites and allosteric effects upon binding. Removing the data analysis bottleneck should enable higher throughput analyses. To illustrate the versatility of the methodology described here, we performed an epitope mapping screen of eight different nanobodies directed against the RBD/SD1 domain of the spike glycoprotein from SARS-CoV2. Each nanobody has a different binding affinity target, expressed in terms of an equilibrium dissociation constant (*K*_D_)^23^. Nine hours were required to collect replicate data for the whole set and only 15 min was needed for data processing (**Figure 7**). Most nanobodies generated a clear zone of protection on the antigen apart from V_H_H 06. The weaker perturbation seems to correlate with its relatively low affinity. These nanobodies were the subject of a previous HX-MS analysis^23^, thus providing us the opportunity to evaluate how well the automated analyses matched the efforts of expert analysts. We identified several distinct binding signatures, which agreed quite well with the previously reported work (**Figure S5**).

**Figure 7.**
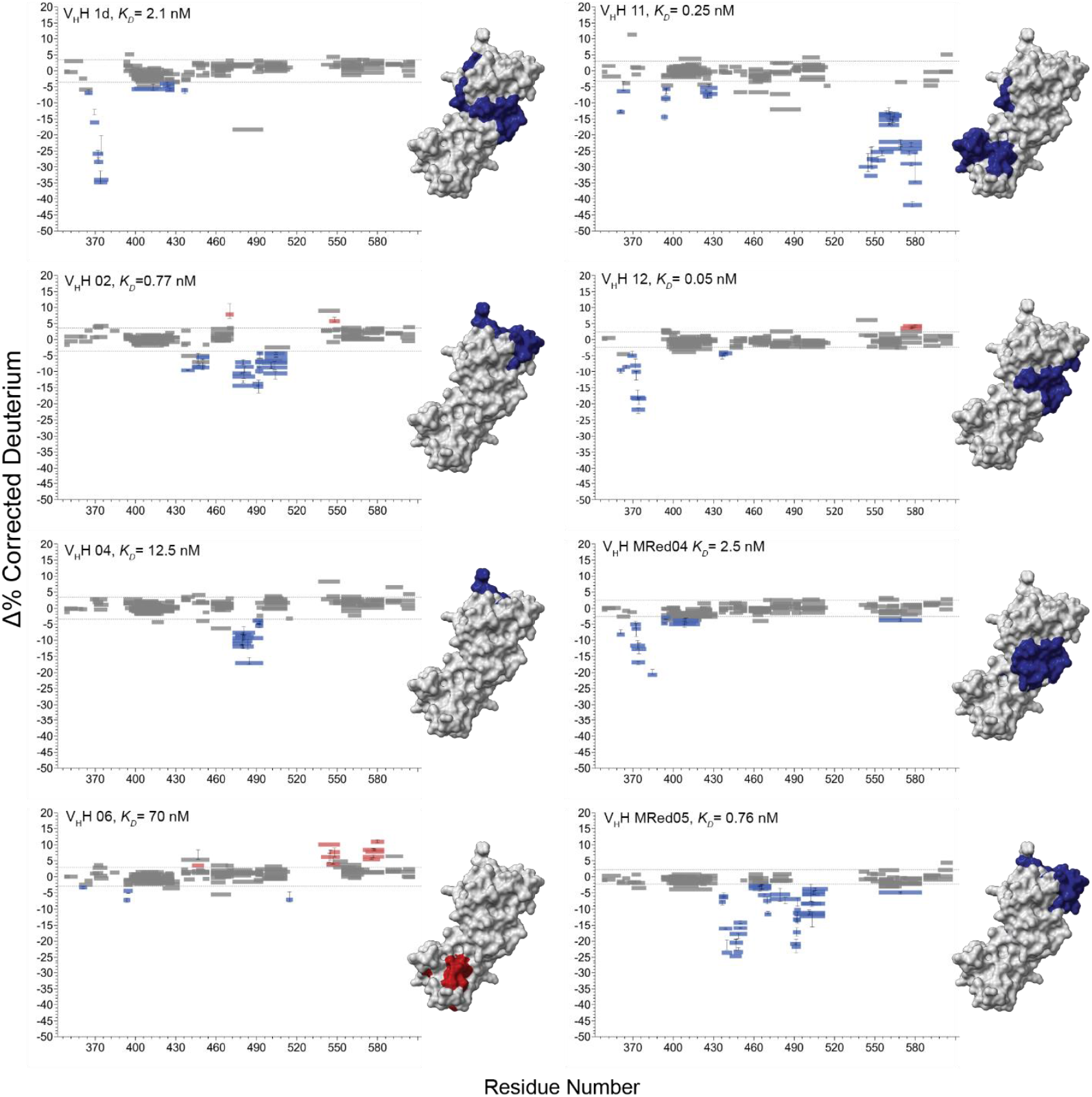
Screening of nanobody binding to the RBD/SD1 (aa319-591) of the spike glycoprotein from SARS-CoV2. Woods plots demonstrating the impact of nanobody binding on deuteration. Nanobody designation and their equilibrium constants, *K*_D_s, for binding to SARS-CoV-2 Wuhan RBD/SD1 are shown^23^. Reductions in deuteration shown in blue, and increases in red, without graduations for the magnitude of change. Perturbations are mapped on 3D structures (PDB 6VXX) in a fixed orientation. Data collected in triplicate.

## Conclusions

HX-MS^2^-based measurements of deuterium exchange were presented over 15 years ago but are only now practical with the advent of improved windowed DIA technologies, and the development of software that can mine higher dimensionality datasets. We demonstrate that standard instrument settings used for DIA-based proteomics applications are suitable for HX-MS^2^, except for the designation of windows and their placement. A reliable protocol begins with an inter-scan method with 20 Th windows and a 10 Th offset, with adaptations based on chromatographic resolution and instrument scan speed. The AutoHX software is easy to use. The user only needs to specify chromatographic peak width and mass tolerances. The minimum number of fragments per peptide and the acceptable precision tolerances for the deuteration calculations are post-processing variables that can be tuned to the quality of the data and the complexity of the protein system.

HX-MS^2^ methods will be more effective on newer mass spectrometers but older and/or slower DIA technologies can still be used to process simpler protein states. These instruments require larger DIA windows to achieve acceptable sampling rates. Even whole mass range fragmentations can be processed by AutoHX (*i*.*e*., MS^E^ on Waters platforms). However, our study shows that noise reduction with smaller windows has a strong positive effect on performance. In summary, the dramatically improved reproducibility, speed and precision of automated HX-MS^2^ analyses are compelling reasons to shift away from conventional HX-MS methods. Further improvements to lab automation will allow the technique to approach high-throughput status and deliver a robust structure-based assay that can be used reliably in a variety of settings, including drug discovery.

## Supporting Information

- Figure S1: Impact of fragment selection on sequence-related metrics. Figure S2: Impact of window overlap on the intra-scan DIA method. Figure S3: Effect of total window size on ion injection times over the course of 50 pmol injections of phosphorylase B. Figure S4: Optimized peptide sequence map for phosphorylase B. Figure S5: Differential deuteration profiles for nanobodies binding to the RBD/SD1 (aa319-591) of the spike protein from SARS CoV2 for two different studies.

## Author contributions

Conceptualization, F.F., D.C.S.; methodology and experiment design, F.F., D.C.S., Mo.H., V.S.; experiments, F.F., Mo.H., Ma.H., M.A.R., J.T., J.G.S., J.H.; project organization, resources and funding acquisition. D.C.S., S.C., R.V.; data analysis and visualization, Mo.H., F.F., V.S., D.A.C., D.C.S.; manuscript writing, D.C.S, F.F., Mo.H.

## Notes

R.V. and Y.S. are employees of Thermo Scientific. S.C. and V.S are employees of Trajan Scientific. AutoHX is now part of Trajan Scientific and Medical’s automation technology platform. M.A.R. and J.T. declare the following competing interests. National Research Council Canada has filed a patent (PCT/IB2022/053756, “Antibodies that bind SARS-CoV-2 spike protein”) that includes the eight nanobodies described here with M.A.R. and J.T. named as inventors. All other authors declare no competing interests.

## Acknowledgements

This work was funded by the Natural Sciences and Engineering Research Council of Canada Discovery Grants RGPIN 2017-04879 to D.C.S. All figures created and/or assembled with BioRender.com, released under a Creative Commons Attribution-NonCommercial-NoDerivs 4.0 International license.

