## Supplementary Data for "Recommendations for Automating Hydrogen/Deuterium Exchange Mass Spectrometry Measurements using Data-Independent Acquisition Methods"

**[#]Authors contributed equally**

**Supporting Information**

### Table of Contents:

1. **Figure S1.** Impact of fragment selection on sequence-related metrics for an HX-MS experiment.

2. **Figure S2.** Impact of window overlap on the intra-scan DIA method.

3. **Figure S3.** Effect of total window size on ion injection times over the course of 100 pmol injections of phosphorylase B.

4. **Figure S4.** Optimized peptide sequence map for phosphorylase B, corresponding to Figure 6 in the main text.

5. **Figure S5.** Differential deuteration profiles arising from a set of nanobodies binding to the RBD/SD1 of the spike protein from SARS CoV2 for two different studies.

### Supporting Information

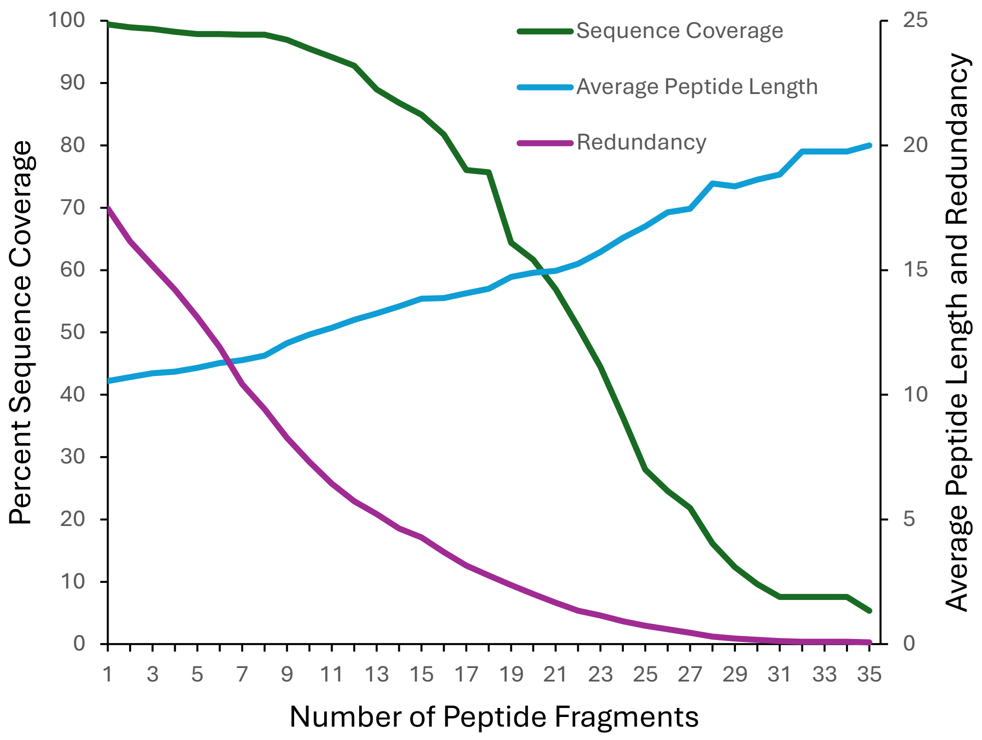


**Figure S1. Impact of fragment selection on sequence-related metrics for an HX-MS experiment.** Plot shows the effect of requiring a progressively higher minimum number of fragments per peptide on percent sequence coverage, average peptide length, and redundancy for phosphorylase B.

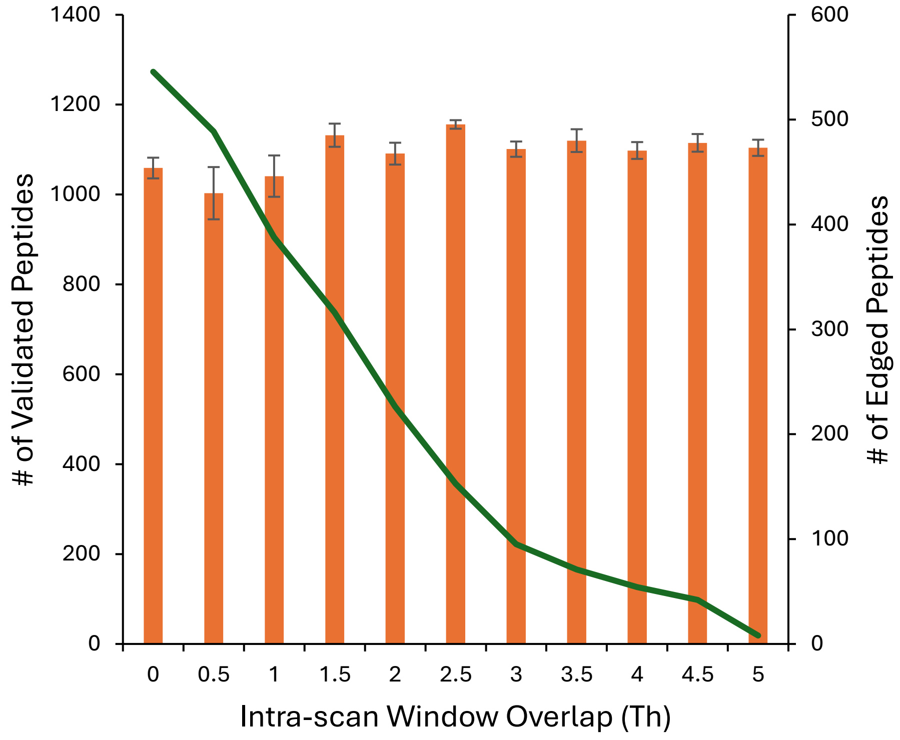


**Figure S2. Impact of window overlap on the intra-scan DIA method.** Plot shows the effect of increasing the overlap on a default window of 10 Th on the # of valid peptides recalled (orange bars) and the number of edged peptides (green line), for a 50% $D_2O$ labeling experiment.

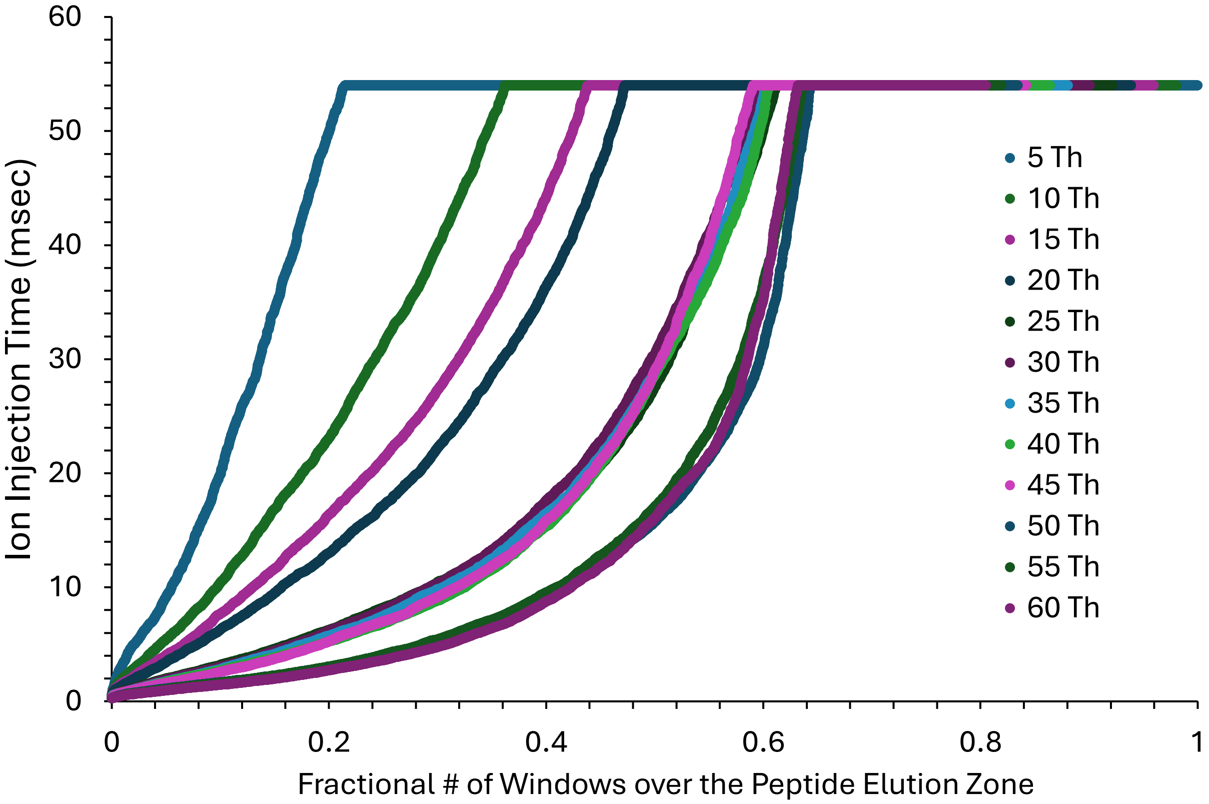


**Figure S3. Effect of window size on ion injection times over the course of 100 pmol injections of phosphorylase B.** Cumulative distribution functions for each overlap tested for the intra-scan method (2 Th default overlap plus the indicated window), expressed as a function of the fractional number of windows sampled across the elution range of the LC-MS/MS runs. Data measured using an AGC setting of 300%. For example, at the recommended 20 Th window, 53% of the windows applied reached the maximum fill time (set at 54 msec).

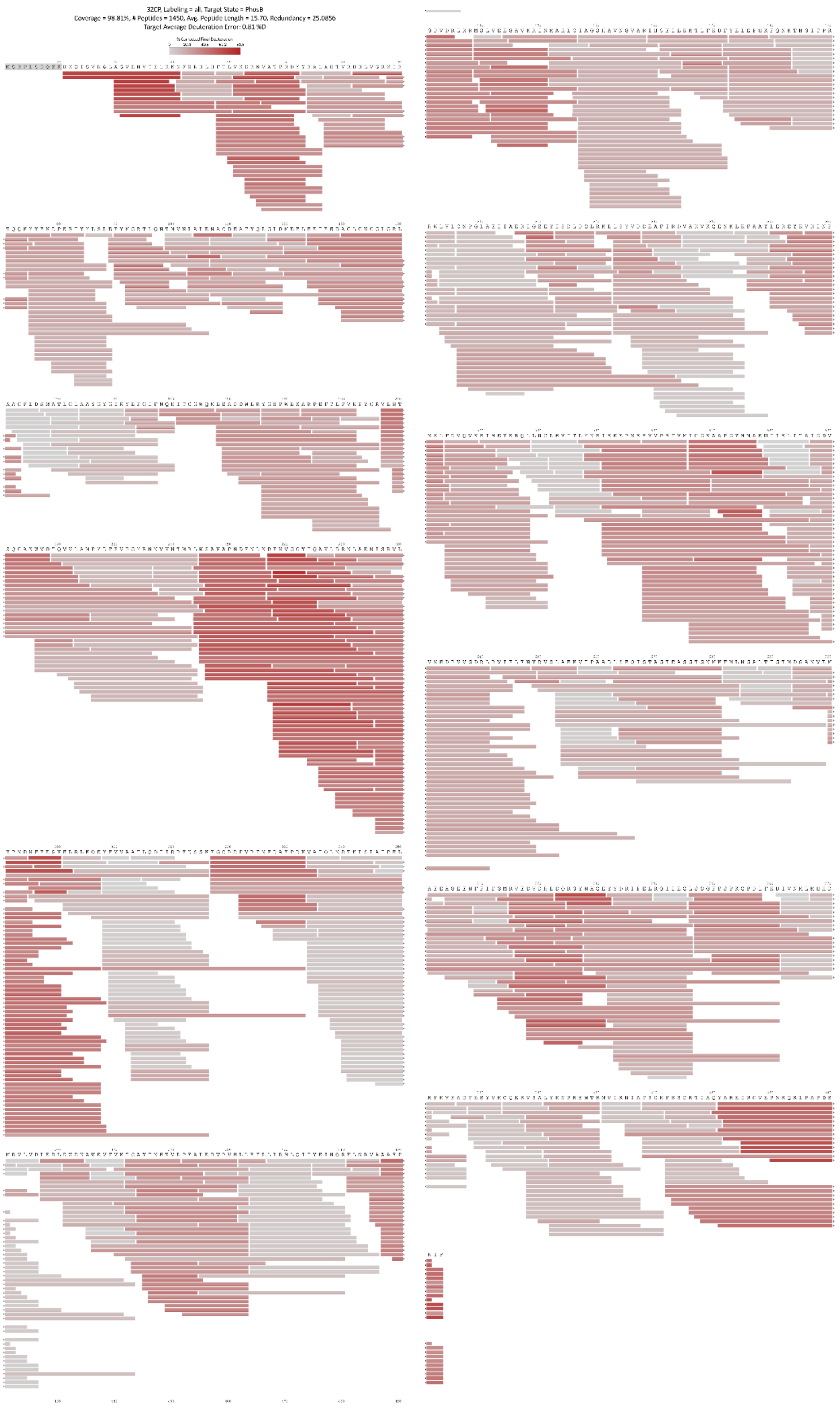


**Figure S4. Optimized peptide sequence map for phosphorylase B, corresponding to Figure 6 in the main text.**

#### Supporting Information

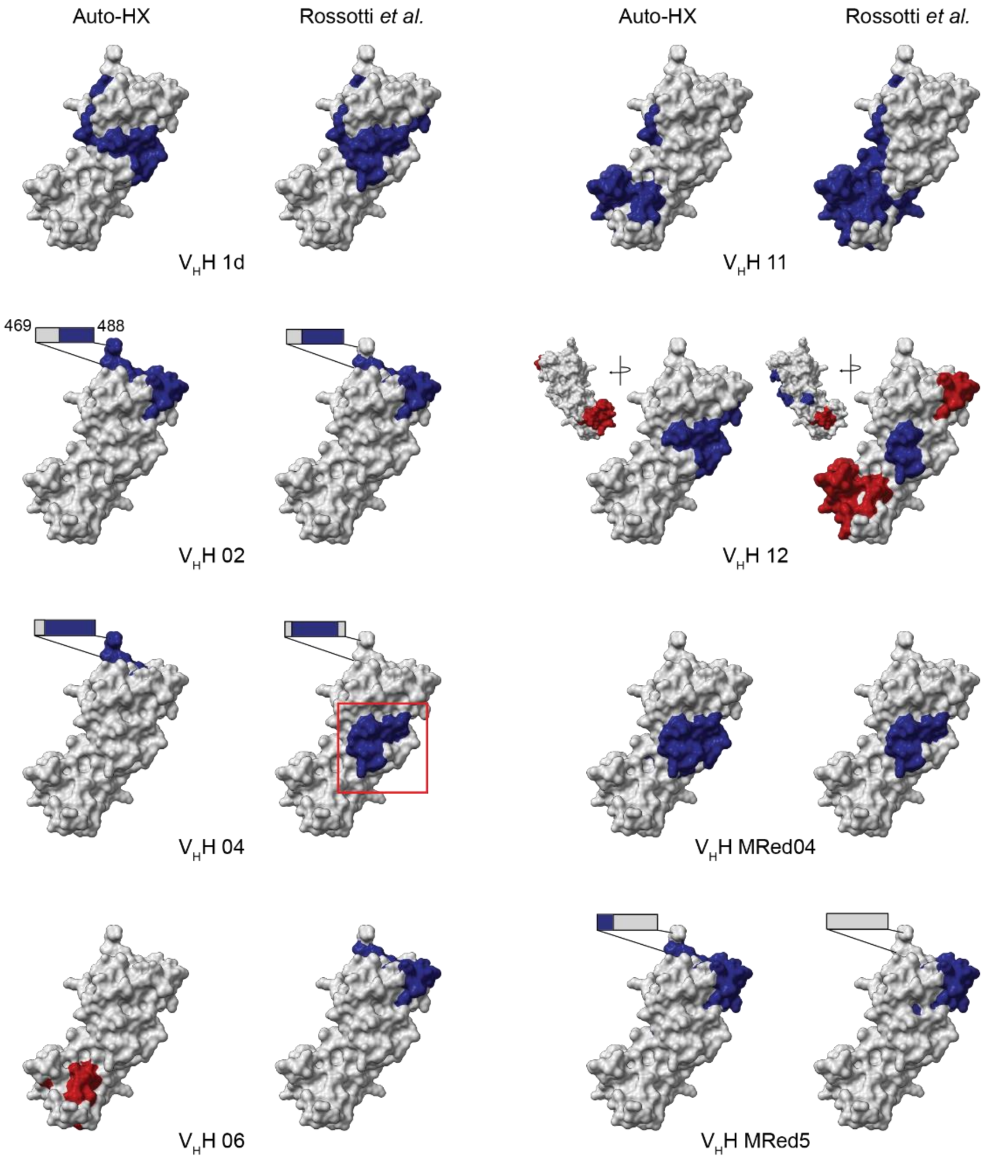


**Figure S5. Differential deuteration profiles arising from a set of nanobodies binding to the RBD of the stalk protein from SARS CoV2 for two different studies.** Comparison of the results from this work and from the Rossotti *et al.* study (reference 23). Reductions in deuteration shown in blue, and increases in red, without graduations for the magnitude of change. Perturbations are mapped on 3D structures (PDB 6VXX) in a fixed orientation. HX results mapped to nominally disordered regions are highlighted using rectangles adjacent to the closest structural region. There is generally good agreement between these results. Note that the difference in $V_HH$ 04 (red box) represents a statistically significant but small change, as do the differences between $V_HH$ 06 and $V_HH$ 12.
